## Supplementary material for "Tracking the popularity and outcomes of all bioRxiv preprints": database schema description

### Supplementary file 1: Database schema

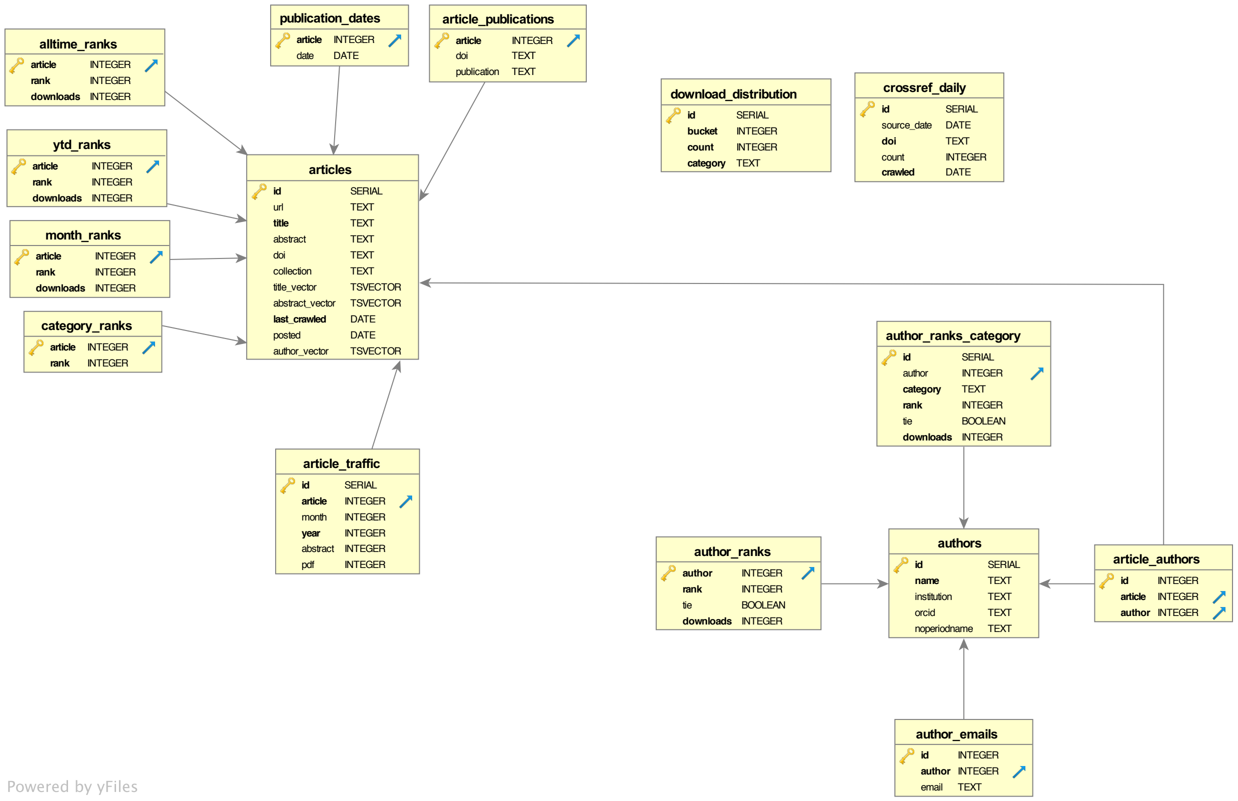


*A graphical illustration of the connections between 15 of the database tables, built using DbVisualizer (https://www.dbvis.com). Each yellow box represents a table in the database, and the entries in each box represent the fields in that table and the type of data stored in each one. The arrows indicate foreign key relationships—references to other tables. For example, the "author_emails" table has an "author" field that has a foreign key relationship with the "authors" table, indicated with an arrow between the two boxes: Any entries in that "author" field will correspond to a value from the "id" field in the "authors" table. (One table, "author_translations," is excluded here. See the table description below for why this is not a table users will need.)*

Rxivist stores bioRxiv data in 16 tables in a PostgreSQL database. It is possible these fields may change in the future to add more data and solidify the schema (more specific constraints, established foreign key relationships, improved primary keys, etc.); changes will be published on rxivist.org and as addendums in the data dumps. Postgres also uses organizational tools called "schemas," namespace-like entities that hold tables. There are two schemas currently available: "prod" contains the most up-to-date information pulled by the Rxivist web crawler. The "paper" schema stores a static copy of the data used for this manuscript, and contains data on all papers posted through the end of November 2018. Currently, the table definitions for both the "prod" and "paper" schemas are identical.

There are several tables that contain data that do not contain the most current data: Any tables in the database with the "_working" suffix have identical fields to the version of the table without that suffix, but contain the previous version of that table. Temporary tables are used to ensure new rankings are loaded properly before "activating" them by renaming the temporary table to the one used by the API, and if an error occurs, the previous rankings can be restored. All tables are listed below in alphabetical order except for "articles" and "authors," which contain the entities central to the data model. See supplementary file *figures.md* for examples of queries that may be helpful.

**Table: articles**

Each entry represents an individual preprint.

- id (serial, primary key) – An arbitrary integer assigned to each preprint when it is first recorded by Rxivist. These values start at 386.
- url (text, unique) – The web address of the preprint's main page on biorxiv.org. Revisions of a paper are given new URLs; this field stores the most recently observed version.
- title (text) – The title of the preprint. This field is updated for revisions.
- abstract (text) – The abstract for the preprint, as specified on bioRxiv. This field is updated for revisions, but abstracts are recorded in a separate step from the one in which preprints and revisions are recorded, so this field may occasionally be null.
- doi (text, unique) – The Digital Object Identifier assigned to the preprint. BioRxiv does *not* issue new DOI numbers for revisions.
- collection (text) – The bioRxiv "collection" in which this preprint has been posted. Because a preprint's collection can only be determined by its appearance in bioRxiv's chronological listing of papers for each category, there is sometimes a delay in which this field will be null.
- title_vector (tsvector) – A sorted list of distinct lexemes found in the preprint's title, as determined by Postgres. Used to facilitate text search. Updated for revisions.
- abstract_vector (tsvector) – A sorted list of distinct lexemes found in the preprint's abstract, as determined by Postgres. Used to facilitate text search. Updated for revisions.
- last_crawled (date) – A date indicating the last time the web crawler checked bioRxiv for updated traffic metrics for the preprint.

- posted (date) – The date on which the preprint first appeared on bioRxiv. This field is *not* updated for revisions.
- author_vector (tsvector) – A sorted list of distinct lexemes found in the preprint's list of authors, as determined by Postgres. Used to facilitate text search. Updated for revisions.

**Table: authors**

Each entry represents an individual preprint author.

- id (serial, primary key) – An arbitrary integer assigned to each author when they are first recorded by Rxivist. These values start at 200,058.
- name (text) – The full name of the author as it was first recorded by Rxivist. Because the back catalog of preprints was not recorded in chronological order and the author name string comparison process changed over time, this name is not necessarily the one attached to an author's earliest preprint.
- institution (text) – The institutional affiliation associated with the author on their most recent preprint. This is updated for revisions.
- orcid (text, unique) – The ORCiD unique identifier associated with the author. This field is populated only if the author has listed the ID on at least one of their preprints.
- noperiodname (text) – The author's full name stripped of all full stops. This field is used to more quickly search for authors without accounting for the punctuation that is applied most inconsistently.

**Table: alltime_ranks**
Each entry stores download ranking information for a single preprint. This table is emptied and replaced every time rankings are calculated.

- article (integer, primary key) – The ID of a preprint.
- rank (integer) – The ordinal position of the preprint in the list of all indexed preprints, organized in descending order according to the "downloads" field. (These rankings do *not* account for ties, so two preprints with the same download count will receive sequential ranks.)
- downloads (integer) – The total overall downloads for the preprint, as of the last ranking calculation. Rankings are not necessarily re-calculated every time traffic data is updated, so this number may be lower than the preprint's total downloads as recorded in the "article_traffic" table, from which this number is calculated.

**Table: article_authors**
Associative table recording the many-to-many relationship between preprints and authors.

- id (serial, primary key) – An arbitrary integer assigned to each association.
- article (integer) – The ID of the preprint being associated.
- author (integer) – The ID of the author being associated.

**Table: article_publications**
Each entry contains the publication information about a single preprint; only preprints that have been published elsewhere have entries.

- article (integer, primary key) – The ID of a preprint.
- doi (text) – The new Digital Object Identifier associated with the published version of the preprint.
- publication (text) – The name of the journal in which the preprint was published, as reported by bioRxiv.

**Table: article_traffic**
Each entry contains bioRxiv traffic data for a single preprint in a single month. Metrics are as reported by bioRxiv, in each preprint's "Metrics" page, in the "Article Usage" section.

- id (serial, primary key) – An arbitrary integer assigned to each entry.
- article (integer) – The ID of a preprint.
- month (integer) – The (1-indexed) month in which the metrics were tallied.
- year (integer) – The four-digit year in which the metrics were tallied.
- abstract (integer) – The number of times the preprint's abstract was viewed on bioRxiv in the specified month.
- pdf (integer) – The number of times the preprint's full-text PDF was downloaded in the specified month.

**Table: author_emails**
Each entry associates an author with an email address. Note there is no requirement that the email be unique, so multiple authors may be associated with the same email address, albeit in a denormalized format. For now, logic in the web crawler avoids associating an author with the same email address multiple times.

- id (serial, primary key) – An arbitrary integer assigned to each entry.
- author (integer) – The ID of an author.
- email (text) – An email address that was associated with that author on the bioRxiv page of one of their preprints.

**Table: author_ranks**
Each entry stores download ranking information for a single author. This table is emptied and replaced every time rankings are calculated.

- author (integer, primary key) – The ID of an author.
- rank (integer) – The ordinal position of the author in the list of all authors, organized in descending order according to the combined all-time downloads of their preprints. (Unlike the alltime_ranks table, these rankings *do* account for ties, so two authors with the same download count will receive identical values in this field.)
- tie (boolean) – A flag indicating whether the author is tied with one or more other authors at the same rank.
- downloads (integer) – The total overall downloads for all preprints associated with the author, as of the last ranking calculation. Rankings are not necessarily re-calculated every time traffic data is updated, so this number may be lower than the total downloads as recorded in the "article_traffic" table, from which this number is calculated.

**Table: author_ranks_category**

Each entry stores download ranking information for a single author in a single bioRxiv category. Only authors with more than zero downloads in a category will have an entry for that category. This table is emptied and replaced every time rankings are calculated.

- id (serial, primary key) – An arbitrary integer assigned to each entry.
- author (integer) – The ID of an author.
- category (text) – Which of the 27 bioRxiv categories was used to limit the download count.
- rank (integer) – The ordinal position of the author in the list of all authors, organized in descending order according to the combined all-time downloads of their preprints in the specified category. (Unlike the alltime_ranks table, these rankings *do* account for ties, so two authors with the same download count will receive identical values in this field.)
- tie (boolean) – A flag indicating whether the author is tied with one or more other authors at the same rank.
- downloads (integer) – The total overall downloads for all preprints in the specified category that are associated with the author, as of the last ranking calculation. Rankings are not necessarily re-calculated every time traffic data is updated, so this number may be lower than the total downloads as recorded in the "article_traffic" table, from which this number is calculated.

**Table: author_translations**

A deprecated table used to specify redirects to search engines that indexed outdated ID numbers for authors that were later modified. Of no practical use elsewhere, but included here because the table is included in the "paper" schema and the database snapshot accompanying this manuscript.

- old (integer, primary key) – An author ID that may have been indexed by search engines but has since changed.
- new (integer) – The new author ID associated with the same individual.

**Table: category_ranks**

Each entry stores download ranking information for a single preprint in the category to which it was posted. (Since each preprint is posted only to a single category, the category itself is not included here.) This table is emptied and replaced every time rankings are calculated.

- article (integer, primary key) – The ID of a preprint.
- rank (integer) – The ordinal position of the preprint in the list of all preprints in the same category, organized in descending order according to all-time download count. (These rankings do *not* account for ties, so two preprints with the same download count will receive sequential ranks.)

**Table: crossref_daily**

Each entry records Crossref social media information for a single preprint on a single day. Note that there are many entries in this table that do *not* reference a preprint; to avoid throwing away data for preprints that have not yet been indexed by Rxivist, the web crawler saves tweet counts for all results with a DOI matching the bioRxiv DOI prefix (10.1101). However, that prefix is shared by other organizations, so some entries are irrelevant external references.

- id (serial, primary key) – An arbitrary integer assigned to each entry.
- source_date (date) – The date for which the data was collected. (For example, a date of "5 Dec 2017" would indicate events reported by Crossref on that date, regardless of when Rxivist itself actually recorded it.)
- doi (text) – The DOI of an entity that may be a bioRxiv preprint.
- count (integer) – The total number of Twitter posts observed on the specified date that referenced the specified DOI.
- crawled (date) – The date on which this information was retrieved from Crossref.

**Table: download_distribution**

Each entry indicates a bin in a histogram used to measure the distribution of all-time downloads for either authors or preprints. Preprints are recorded in the category "alltime"; authors are in the category "authors". Though it doesn't conform well to the schema, this table is also used to record the mean and median download counts for preprints for authors, preprints, and each of the 27 bioRxiv categories. The "category" field for these entries is the category name plus either "_median" or "_mean" at the end. The "bin" field for these entries is "0" and the "count" field is the value. These values are used for visualizations on the Rxivist website.

- id (serial, primary key) – An arbitrary integer assigned to each entry.
- bucket (integer) – The maximum number of downloads for this bin.
- count (integer) – The number of entities with a total download count that falls within the limits of this bin.

- category (text) – Which histogram this bin belongs to.

**Table: month_ranks**
Each entry stores download ranking information for a single preprint based on download data going back to the beginning of the previous month. This table is emptied and replaced every time rankings are calculated.

- article (integer, primary key) – The ID of a preprint.
- rank (integer) – The ordinal position of the preprint in the list of all indexed preprints, organized in descending order according to the "downloads" field. (These rankings do *not* account for ties, so two preprints with the same download count will receive sequential ranks.)
- downloads (integer) – The total downloads for the preprint since the beginning of the previous month, as of the last ranking calculation. Rankings are not necessarily re-calculated every time traffic data is updated, so this number may be lower than the total as recorded in the "article_traffic" table, from which this number is calculated. However, ranks for all preprints are calculated at the same time, so the timeframe covered by this field will be the same across the whole table.

**Table: publication_dates**
Each entry stores publication information for a single preprint. Most (but *not* all) preprints that are recorded in the "article_publications" table have a corresponding entry in this table. This is the only table present in the "paper" schema but not the "prod" schema and is not maintained.

- article (integer, primary key) – The ID of a preprint.
- date (date) – The date on which the preprint was published in a journal, based on data from Crossref.

**Table: ytd_ranks**
Each entry stores download ranking information for a single preprint based on download data going back to the beginning of the current year. This table is emptied and replaced every time rankings are calculated.

- article (integer, primary key) – The ID of a preprint.
- rank (integer) – The ordinal position of the preprint in the list of all indexed preprints, organized in descending order according to the "downloads" field. (These rankings do *not* account for ties, so two preprints with the same download count will receive sequential ranks.)
- downloads (integer) – The total downloads for the preprint since the beginning of the current year, as of the last ranking calculation. Rankings are not necessarily re-calculated every time traffic data is updated, so this number may be lower

than the total as recorded in the "article_traffic" table, from which this number is calculated. However, ranks for all preprints are calculated at the same time, so the timeframe covered by this field will be the same across the whole table.
